# The Direct Subjective Refraction: Unsupervised measurements of the subjective refraction using defocus waves

**DOI:** 10.1101/2021.12.04.471123

**Authors:** Victor Rodriguez -Lopez, Alfonso Hernandez-Poyatos, Carlos Dorronsoro

## Abstract

We present the Direct Subjective Refraction (DSR), a new subjective refraction method, and validate it vs the Traditional Subjective Refraction (TSR) and an unsupervised version of it (UTSR). We project an optotunable lens onto the eye to create Temporal Defocus Waves produces flicker and chromatic distortions, minimum when the mean optical power of the wave matches the spherical equivalent of the eye. 25 subjects performed the DSR visual and UTSR tasks without supervision. DSR is more repeatable than TSR and UTSR (standard deviations ±0.17D, ±0.28, and ±0.47D). The time per repetition of DSR is only 39s (almost 6 min for TSR). Cyclopegia severely affects UTSR, but not DSR, confirming that the DSR task de-activates the accommodative system. DSR is a new method to obtain the spherical equivalent that does not requires supervision and overpasses existing subjective methods in terms of accuracy, precision, and measurement time.

## INTRODUCTION

The gold standard to assess the refractive error of an eye is the subjective refraction [1,2]. This procedure is probably the most frequent procedure performed in eye care clinics, as it is not only performed on patients candidate to an optical correction, but also as a method to obtain basic information about the state of the patient’s eyes before many other ophthalmic procedures. Therefore, in most eye care clinics, subjective refraction is performed on most patients.

The purpose of subjective refraction is to find the most suitable combination of lenses that compensates for the refractive error of an eye, producing the maximum visual acuity. In the final steps, practitioners iteratively ask the patient about the perceived blur of a visual test (usually letters), trying different lenses and aiming to minimize said blur. Due to the natural blur of the eye, the optical aberrations, and the resultant depth of focus, the blur-detection task is in fact a blur-preference task that is challenging both for the patients and for the practitioners, often substituted with a letter identification task, that has the same drawbacks. The responses are often dubious, and the practitioners must guide the patients and interpret their subjective feedback.

Traditionally, objective refraction instruments (retinoscopes, autorefractors) are used to provide a first approximation to the patient’s refraction, which has to be subsequently refined [1]. The subjective refraction procedure thus begins by adding more positive power, usually +1.00 diopter (D), to the starting point to induce myopic defocus, and then reducing said addition in steps of 0.25D until the best visual acuity is reached. This method, known as fogging, is used to relax the eye, reducing the impact of accommodation in the measurement. Other subjective refraction techniques, such as the duochrome test, can refine the result. The duochrome test consists of black letters displayed on red and green backgrounds, and takes advantage of the longitudinal chromatic aberration of the eye (different wavelengths focus at different points with respect to the retina). The dioptric difference between the red and green component is about 0.50D and the subject’s task is to indicate which background produces better-focused letters, providing the practitioner with a useful hint about the direction of the residual refraction, and guiding a few more subjective refraction iterations.

Different technologies have been used to support subjective refraction procedures. The most common, trial frames, manual phoropters, and digital phoropters, provide high correlations and non-significant differences. [3]

Subjective refraction is time-consuming. For each patient, it entails the full dedication of a well-trained eye care professional. For many eye care practitioners, subjective refraction represents a considerable fraction of every workday, estimated to be about 6 minutes per eye [3–5].

The subjective nature of the subjective refraction implies deviations in the measurement result. The standard deviation across repetitions performed by the same optometrist evaluating the same eye -intraoptometrist variability-has been reported to be, in the spherical component around ±0.25D [6–8]. The standard deviation across repetitions performed by different optometrists evaluating the same eye -interoptometrist variability-, has been reported to be slightly higher (in spherical component) than the intraoptometrist variability, ±0.28D [7,9–12]. It has been reported that this variability can cause headaches and distortions [13], intolerances [14], and differences in perceptual preference without reduction of visual acuity [15], highlighting the importance reducing the variability.

In an attempt to reduce the measurement time and facilitate both the task of the patient and the complexity of the procedure followed by the practitioner, many objective refraction technologies, without the need for subjective responses from the patient, have been improved throughout the years.

Autorefractors provide a direct measurement of the refractive error of the patient. Classic autorefractors project optical objects (typically dots or rings) onto the eye and obtain the refractive state of the eye from the analysis of the retinal image (typically the size or the blur of the image), using different technologies such as infrared lasers, LEDs, superluminescent diodes, Badal systems, CCD or CMOS cameras [16]. Many of them also incorporate fogging algorithms. Moreover, new technologies based on wavefront analysis have emerged over the years [17]. Several studies have reported very high repeatability of autorefractometers, with a standard deviation of repeated measurements ranging from ±0.12 to ±0.37D [6,7,18,19].

Other studies have reported significant differences in spherical equivalent between autorefractometers and subjective refraction. Thibos et al. [17] studied the precision of 33 different metrics derived from the wavefront aberration map of the eye, to predict the spherical equivalent of the refractive error. They found mean absolute differences on the spherical equivalent, compared with the subjective refraction, ranging from ±0.25 to ±0.48D, giving birth, in 2004, to wavefront autorefractometry, a discipline that has provided several autorefraction technologies over the years [7,10,18,20–24]. Modern autorefractometers provide mean absolute differences ranging from ±0.55 to ±0.24D, depending on the autorefractometer [20,25]. A recent review [26] analyzed four portable autorefractors and reported that QuickSee provides the lowest mean absolute difference (±0.21D) [27]. Many other studies have compared subjective refraction with different types of autorefractors, reporting similar differences [10,19,21–24]. To summarize, both classic autorefraction and wavefront-based autorefraction provide high repeatability, with wavefront autorefractors providing better predictions of the spherical component of the subjective refraction (mean absolute deviations between ±0.25 and ±0.50D). Other objective tools, such as deep-learning algorithms [28], have been used to predict the spherical refractive error from retinal fundus images, but their outcomes are still far from other objective techniques (±0.91D with respect to the subjective refraction).

Interestingly, the outcome of subjective refraction is better visually accepted than the outcome of autorefraction [9,29] with higher visual acuity reported with subjective refraction than with autorefraction [30,31].

Some recent developments advance toward self-refraction and unsupervised subjective refraction. Sheedy et al. [7] compared an automatic subjective refraction system, the Topcon BV-1000 with computerized forced-choice questions, with autorefraction and with the traditional subjective refraction. The virtual subjective refraction [5] was proposed as a semi-automatic subjective refraction method that combines wavefront-based objective refraction providing a starting point, and an algorithm driving a virtual-reality system and processing the responses of the subject. The EYER method [4] combines a wavefront autorefractor and a phoropter to, based on the answers of the subjects to specific questions, change the lenses of a phoropter and obtain the refractive error subjectively. Leube et al. [32] tested an unsupervised self-refraction procedure based on the shift of a pair of Alvarez lenses to compensate for the spherical refractive error. Rotation of the lenses also allows the compensation of astigmatism. Wisse et al. [33] tested a web-based tool (different letter tests and algorithms) to measure the refractive error. These methods report mean absolute difference with respect to the subjective refraction between ±0.41 and ±0.21D. Furthermore, Elliot [2] suggested questionnaires based on patient satisfaction and preference [34]. The main inconvenience is that this procedure is time-consuming because it implies testing different corrections for several days until the patient feels satisfied with the prescription

As already mentioned, accommodation is an important issue in refractive error evaluation, common to objective and subjective refraction techniques, especially in young populations [21]. Different strategies aim at reducing its impact. Even with the fogging method, traditionally used in subjective refraction, and later incorporated into certain objective methods, mild or higher hyperopias are often missed. Cycloplegic drugs can null the influence of accommodation, although they produce other unwanted effects such as pupil dilation, visual discomfort, or photophobia. Zadnik et al. [6] reported a higher standard deviation (lower repeatability) in cycloplegic subjective refraction versus the non-cycloplegic subjective refraction (±0.48 vs ±0.32D). Additionally, Choong et al. [35] and Rauscher et al. [36] reported, in a population of 117 and 201 children, respectively, that cyclopegic autorefraction was more accurate than non-cycloplegic, that actually gives more myopic outcomes (∼0.5D). Overcorrection of myopia or undercorrection of hyperopia, mainly provoked by accommodation, can produce asthenopia and headache [37]. Those symptoms could be eliminated with a better estimation, insensitive to accommodation, of the refractive error.

The current trend is to direct the evaluation of the refractive error toward automatic unsupervised methods, that minimize: 1) miscommunications between the clinician and the patient; 2) the measurement variability; 3) the measurement time; 4) the influence of accommodation (without the need of cyclopegic drugs). Subjective methods are likely to achieve results that are closer to the traditional subjective refraction, nowadays still considered the universal gold standard in the evaluation of refractive error.

## THE DIRECT SUBJECTIVE REFRACTION CONCEPT

The Direct Subjective Refraction (DSR), a new concept proposed in this study, is a subjective method to obtain the spherical equivalent of an eye, i.e., the optical power needed to compensate for the spherical refractive error. It is based on inducing rapid and periodic temporal changes in the optical power to an eye (Temporal Defocus Waves; TDWs) and therefore producing periodic temporal changes in the focus state of the retinal image while maintaining its position and magnification [38]. These fast periodic changes in defocus produce periodic changes in retinal blur and the visual perception of flicker in the image, which is minimum when the mean optical power of the TDW corresponds to the spherical equivalent of the subjective refraction of the eye. The flicker increases as the mean optical power of the TDW moves away from the spherical equivalent becoming more and more myopic or hyperopic.

In this method, the stimulus is made of different chromatic components, for example blue and red monochromatic components and combinations of them. Due to the Longitudinal Chromatic Aberration (LCA) of the eye, each monochromatic component is focused at a different axial position relative to the retina. The TDW interacts with the LCA producing color distortions in the stimulus and differences in the flicker of the different chromatic components. These color distortions are minimum, again, when the mean optical power of the TDW matches the spherical equivalent of the eye. Interestingly, the color distortions are different at both sides of the focus.

The task of the observer in the DSR method is to minimize (1) the flicker in the stimulus and (2) the color distortions. Both perceptual effects are dynamic, concurrent, and very apparent to the observer. As a result, the two perceptual effects used in the minimization task reinforce each other to converge to a common focus, making the task easy for the observer. Around the focus, flicker and color distortions are image features perceptually stronger than the static blur commonly used to guide the traditional subjective refraction. In other words, the dual minimization task used in DSR is more sensitive than the one used in the traditional subjective refraction (blur reduction) and less affected by accommodation. In this study, we will demonstrate that the results obtained with the DSR method are more robust, precise, and direct (providing faster and more repeatable measurements) than those obtained with the traditional method.

### Working principle

Figure 1 describes the working principle of DSR, based on the dynamic interactions between the LCA of the eye and the fast temporal variations of optical power induced by the TDW (Figures 1A, 1B, and 1C). The through-focus retinal images of the edges of a stimulus are very different across chromatic components due to LCA (as an example: red monochromatic edge through focus in Figure 1D; blue monochromatic edge in Figure 1E; magenta bi-chromatic edge -red plus blue-in Figure 1F). The periodic alternation in defocus generated by the TDW produces quick variations in blur that are perceived as periodic luminance changes, i.e., flicker. At a given defocus, the blur is different for each chromatic component, and the different spread of light produces energy unbalances, changing the color around the edges of the stimulus (Figures 1G and 1H). The observer subjectively selects their refraction by adjusting the mean value of the TDW (i.e. their refractive state) until those perceptual effects -flicker and color artifacts-are reduced to a minimum.

**Fig. 1.**
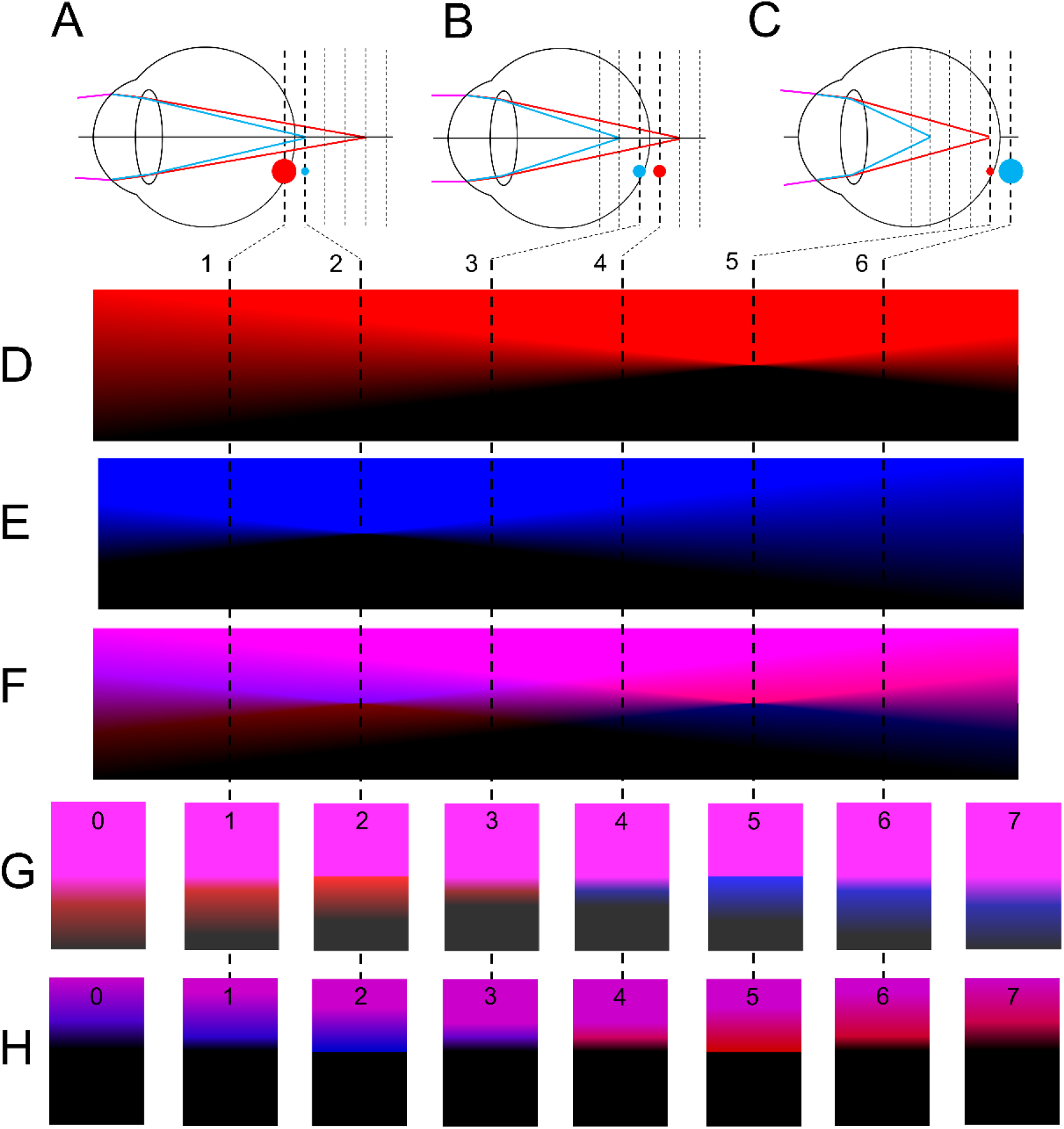
Working principle of the Direct Subjective Refraction. The perception of the stimulus depends on the mean optical power of the TDW, which changes the plane in focus in the retina. **A**. Schematic representation of an eye in a hyperopic state for a given mean value of the TDW. Six optical planes are represented with dashed lines (1 to 6). The two optical powers of the TDW are represented with bold dashed lines: one of them, plane 2, corresponds to the blue focus of the eye and the other one, plane 1, is defocused. **B**. Eye in an emmetropic state with respect to the TDW. It could be the same eye, but with more mean optical power in the TDW. The two optical powers of the TDW correspond to planes 3 and 4, very close and at either side of the retina, i.e., similarly defocused. **C**. Eye in a myopic state with respect to the TDW. It could be the same eye, but with even more mean optical power in the TDW. The two optical powers of the TDW correspond to planes 5, red focus of the eye, and 6, defocused. **D**. Representation of the through-focus blur, induced by defocus, for a red edge. Only plane 5 is in focus. **E**. Through-focus blur for a blue edge, with plane 2 in focus. **F**. Through-focus blur for a magenta edge (red plus blue). The different defocus in the blue and red components induce color distortions that are different in the hyperopic and myopic sides of the retina, and in the bright and dark sides of the edges. **G**. Images corresponding to planes 1 to 6 (and also to additional planes 0 and 7) of a magenta edge with high contrast and brightness, representing the vision of an eye not completely adapted to a bright display. The observers perceive color distortions in the dark side of the edges: reddish tint in the hyperopic side of the retina and blueish tint on the myopic side. **H**. Same images, but with lower contrast and brightness, corresponding to an eye not completely adapted to a dim display. Color distortions are now better perceived in the bright side of the edges: now the tint is blueish in the hyperopic side of the retina, and reddish in the myopic side. In this figure, for illustration purposes, the amplitude of the TDW was only one-third of the chromatic difference of focus between the blue and the red wavelengths. However, the effect will be magnified by a larger amplitude, producing a bigger change in the image of the edges. See *Methods* section (pilot experiments).

Six representative through-focus planes are considered in Figure 1, numbered 1 to 6, and shown as dashed lines in Figures 1A, 1B, and 1C. The TDW is represented by two bold dashed lines indicating the two planes of alternating focus. Figures 1A, 1B, and 1C represent three different refractive states in which the TDW has different mean optical powers with respect to the retina.

In Figure 1A, one of the alternating optical powers of the TDW corresponds to the blue focus of the eye (plane 2) -where the blue components of the stimulus are sharp-, and the other one places the stimulus in front of the retina (plane 1). In this situation, the best focus of the eye (in between the blue and the red foci) would lay behind the retina. This eye is, therefore, in a hyperopic refractive state. The alternation between planes 1 and 2 induced by the TDW produces (see also Visualization 1A): i) More average blur in red image components than in blue image components; ii) More flicker perception in red image components -where blur is suprathreshold in both planes 1 and 2-than in blue image components - were blur is subthreshold in plane 2 and small in plane 1-, and; iii) A reddish halo within the dark side of magenta edges, and blueish halo within the bright side (Figures 1F, 1G and 1H) in both planes 1 and 2. The reason is that in an edge between magenta and black, the red light is spread more than the blue light. In the dark side of the magenta edge, the additional red produces a reddish halo (Figure 1G, planes 1 and 2), and in the bright side, the missing red produces a blueish halo (Figure 1H, planes 1 and 2).

Figure 1C represents the opposite situation. It could represent the same eye, but with more average optical power in the TDW. One of the optical powers corresponds to the red focus (plane 5), and the other one projects the stimulus behind the retina (plane 6). Because the best focus of the eye would lay in front of the retina (between the blue and the red foci), this eye is in a myopic refractive state. In this other case, the observer experiences (see also Visualization 1C): i) More blur in blue than in red; ii) More flicker in blue than in red, and; iii) A blueish halo within the dark side of the magenta edge, and a reddish halo within the bright side (Figures 1G and 1H, planes 5 and 6).

Figure 1B shows the particular situation in which the eye is in focus (it could represent, again, the same eye). In this perfectly corrected eye is focused in between the blue focus -in front of the retina- and the red focus -behind the retina-. The two optical powers of the TDW correspond to planes 3 and 4, very close and at either side of the best retinal focus. The mean optical power of the TDW matches the retinal plane and therefore the eye represented is in an emmetropic state. Consequently (Visualization 1B): i) Blue and red components have similar amounts of blur; ii) Similar small flicker in blue and red components and; iii) Blue and red light are barely spread and the difference is hardly noticeable. The color distortions in magenta edges disappear with the fast alternation because no color dominates the other.

Blur, flicker, or color artifacts keep increasing for higher amounts of myopia (position 0) or hyperopia (position 7). In summary, when the eye is defocused in a TDW scheme, not only the amount of blur increases as defocus increases, but also, and more noticeably, the amount of flicker and color distortions. Even a slight residual defocus results in an increase of these effects. The color distortion is different at both sides of the focus of the eye (blueish tint of black objects if myopic defocus, reddish if hyperopic; Figure 1G). Therefore, color artifacts not only provide an indication of the amount of defocus, but also an unambiguous cue of the defocus sign. Furthermore, the fast change in defocus produced by the TDW also has a beneficial effect on distracting accommodation. The accommodation system is no longer able to follow the (quick) changes in optical power and the image cannot be focused. Therefore, the unstable nature of the image does not provide a cue to the activation of the accommodation mechanism.

This study proposes Direct Subjective Refraction as a new unsupervised subjective refraction method to overcome some of the limitations of the existing objective and subjective refraction methods. In this work, different stimuli have been designed and tested, in combination with the TDW (*Methods* Section, Figures 3 and 4, Visualization 1), to maximize the perceptual effect and the sensitivity of the dual task (minimization of flicker and color distortions). In this method, the task is so straightforward for the patients that they can perform the minimization routine by themselves. The practitioner intervention is reduced to explaining the perceptual task at the beginning of the test, and supervising the result. This research also explores the limits of an unsupervised version of traditional subjective refraction, expected to result in a faster procedure, but inaccurate.

## METHODS

### Optical System

The active part of the optical system is an optotunable lens, a lens able to change its optical power in response to an electric signal. The optotunable lens used in this study is based on liquid-membrane technology (EL-10-30-TC, Optotune, Switzerland), enabling precise changes in optical power up to 100 Hz [39,40]. The TDW is produced by a SimVis simultaneous vision simulator that uses a 4f-projection optical system to optically conjugate the optotunable lens with the pupil plane of the eye of the observer (Figure 2A). The distance from the eye pupil to the first lens is 45mm. The distance from the optotunable lens to the stimulus is 1 meter. The total distance from the real position of the eye to the screen is 1.25m.

**Fig. 2.**
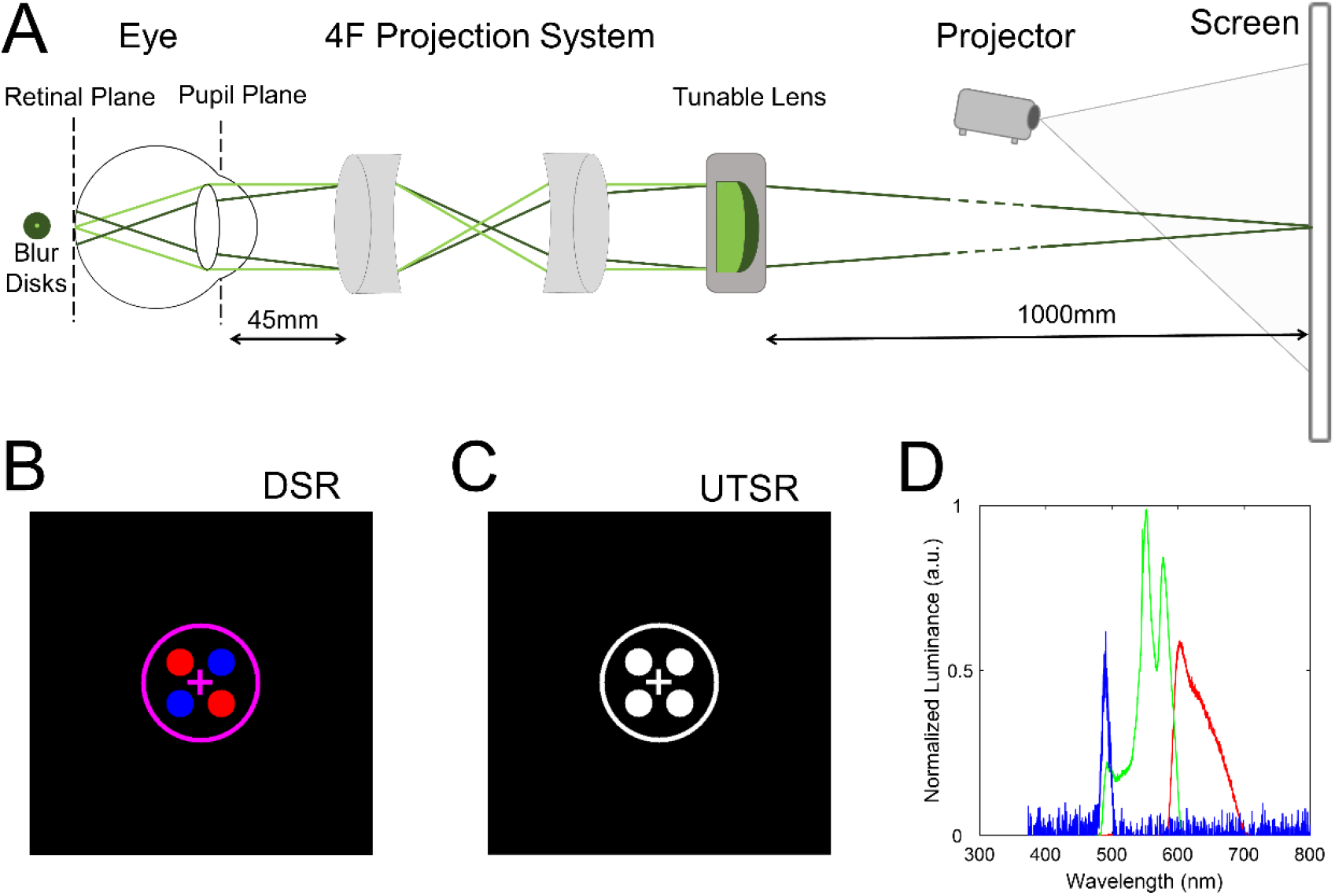
Setup of the study. **A**. Schematic representation of the optical system of the Direct Subjective Refraction. It shows how inducing optical powers with the optotunable lens changes the retinal blur. The 4f optical system projects the optotunable lens on the pupil plane of the eye. In most situations, the stimulus is defocused for the observer, producing a large blur disk on the retina (dark green). In the particular situation when the optotunable lens focuses the stimulus on the retina (light green), the blur disk is minimum. DSR uses fast variations in optical power. With this configuration, the optical power of the optotunable lens produces defocus blur in the image, without changing the position nor the magnification. The sizes and distances displayed are not proportional to the real optical system. **B**. Stimulus used to perform the DSR task. **B**. Stimulus used to perform UTSR task. **C**. Spectral emission of the light source (DLP) for blue, green, and red components. See also Visualization 1 for a simulation of the appearance of the DSR stimulus during the DSR task.

Custom routines were programmed in Matlab (Math-works Inc., Natick, USA) to operate the custom driver based on Arduino electronics (Arduino Nano 3; Arduino, Italy) controlling the optical power of the optotunable lens, and to implement the TDW. Matlab, in combination with Psychtoolbox [41], was also used to design and present the stimuli and to perform the perceptual task.

### Temporal Defocus Wave (TDW)

We performed pilot experiments in experienced subjects to refine the design of the stimuli, and to find the parameters of the TDW that maximized the perceived flicker and the intensity of the chromatic distortions outside the best focus. The stimulus is described in the E*xperiments (3.4 and 3.5)* section and is shown in Figure 2B. The temporal frequency of the TDW was set to 15 Hz, corresponding to the maximum perceptual sensitivity to defocus flicker [42]. The peak-to-peak amplitude of the TDW was set to 0.50D producing a good balance between measurement sensitivity and precision.

With those optimal parameters of the TDW, flicker and chromatic distortions are almost negligible in focus (if the TDW oscillates between planes 3 and 4 in Figure 1), but very apparent out of focus. With smaller amplitudes, the two images of each pair are very similar, and the subject has difficulties perceiving flicker even outside the best focus. With larger amplitudes, the two images are very different in luminance and flicker is very noticeable, even in focus.

### Subjects

Twenty-five subjects, 15 female and 10 male, between the ages of 48 and 23 (29.9±7.3 on average), participated in the study. All participated in Experiment 1, and five of them also in Experiment 2. No color abnormalities were found, tested with Ishihara chromatic test. All subjects had normal visual acuity (VA; ≤0.0 logMAR) wearing their usual correction. Experiments were performed with the room lights switched off.

Objective refraction was measured using an ARK-1 autorefractometer (ARK1, Nidek, Japan). The refractive error (in spherical equivalent) ranged from -6.75 to +1.50D (−1.62±2.32D on average) with a distribution of eight emmetropes (±0.50D or refractive error), fourteen myopes (<-0.50D) and three hyperopes (>+0.50D). Subjects were free to accommodate, except in Experiment 2 where the accommodation was paralyzed using cycloplegic drugs.

A bite bar provided centration stability during the experiments. Fixation was provided by the stimuli. Only the left eye was measured. The right eye was occluded with an eyepatch.

The experimental protocols were approved by the Spanish National Research Council (CSIC) Bioethical Committee and were in compliance with the Declaration of Helsinki. Written informed consent was provided by all subjects.

### Experiment 1

Figure 2B shows the stimulus used to evaluate DSR. This stimulus was designed to intensify the perception of flicker and chromatic artifacts at both sides of the focus.It comprises four circles arranged like the corners of a square, alternatively red (RGB coordinates [1 0 0]) and blue ([0 0 1]), on a black background ([0 0 0]). The diameter of the circles is 1°. They are surrounded by a thin magenta ring ([1 0 1]; 4.7° of visual angle). The stimulus also contains a magenta cross in the center, for fixation.

Visualization 1 shows a computer simulation of the interaction between the DSR stimulus, the TDW, and the LCA of the eye, which produces flicker and chromatic distortions depending on the mean optical power of the TDW. The flicker of the blue dots is preponderant when the mean power of the TDW changes the refractive state of the eye to myopia, while the flicker of red dots becomes more visible in hyperopia. In emmetropia, flicker is minimum and similar for red and blue dots. In the DSR stimulus, the chromatic artifacts appear in the magenta components (the fixation cross and the surrounding ring), which are shifted to blue in myopia and to red in hyperopia.

The DSR task consisted in simultaneously minimizing the two concurrent effects in the image induced by the TDW: the flicker and the chromatic distortions. Subjects increased or decreased the mean power of the TDW using a keyboard, in coarse or fine steps of 0.25D or 0.10D, respectively.

To compare with the DSR, subjects also performed an Unsupervised Traditional Subjective Refraction (UTSR) task, a version of the Traditional Subjective Refraction (TSR) used in clinical practice. The stimuli designed for UTSR (Figure 2C; UTSR) is a black-and-white version of DSR (Figure 2B), in which magenta, blue and red colors are replaced with white ([1 1 1]). The task of the subject was to minimize the blur (defocus) of the stimulus, changing the optical power of the optotunable lens with the keyboard (same steps) until the stimulus was perceived sharp. In this case, the explanation of the task did not require a video presentation and therefore it was faster than in DSR.

The explanation preceding the experiment took about 1.5 minutes for the DSR task and 0.5 minutes for the UTSR task. The time elapsed between the explanation and the conclusion of the unsupervised tasks was recorded in both methods.

Subjects wore their current prescription throughout the experiments (delivered by spectacles or contact lenses). Both in DSR and UTSR, the spherical equivalent was estimated averaging 10 repetitions, each one following a staircase procedure with a different starting point: 5 of them myopic, from -0.20D to -1.00D, and the other 5 hyperopic, from +0.20D to +1.00D. The precision of the method was estimated as the standard deviation across repetitions. As subjects wore their spectacles or contact lenses, both tasks measure residual refraction, i.e., deviations with respect to their current correction. But the results, obtained from different myopic and hyperopic starting points, are representative of arbitrary refractive errors. After the measurements, visual acuity was checked with the spherical equivalent obtained with the DSR task.

The display was a combination of a digital light projector (DLP PJD7820HD, ViewSonic, USA) and a flat white reflecting screen. The distance from the projector to the screen was 0.4 meters, providing a sharp image with high luminance (500 cd/m^2^ if set to white). The spectral emission, plotted in Figure 2D, shows narrow and close R and B components.

### Experiment 2

In Experiment 2, five subjects performed the same procedure and using the same experimental setup as in Experiment 1, but with the accommodative response paralyzed after the instillation of cyclopegic drugs (tropicamide 1%). Measurements began 10 minutes after the instillation of the third dose.

### Statistical Analysis

To analyze the statistical significance of the differences between the results of the different experiments, we used Wilcoxon signed-rank tests to compare the results of Experiment 1 vs 2. Additionally, to compare groups with different refractive errors (myopes, hyperopes, and emmetropes), we also used a Mann-Whitney U-test for different sample sizes.

To evaluate the agreement between DSR and UTSR tasks versus the TSR, we used Bland-Altman plots. The statistical level to achieve statistical significance was set to 5% (p=0.05). For each subject, we considered each repetition of the DSR and UTSR task as a different measurement of residual refraction. Additionally, paired t-tests and correlation coefficients were also used to compare between myopic and hyperopic starting points in the DSR and UTSR tasks. Matlab (Math-works Inc., Natick, USA) was used to perform the analysis.

## RESULTS

Figure 3 depicts a few representative examples of the measurements performed. Each panel shows the progress along trials (staircase) of every repetition for Subject S#1, Experiments 1 and 2 and both tasks, DSR and UTSR. Blue lines represent hyperopic starting points and red lines myopic starting points. The X-axis represents the trial number. The Y-axis represents the mean optical power of the TDW, in diopters, for the Direct Subjective Refraction (DSR; examples in panels A and C) and the optical power, in diopters, for the Unsupervised Traditional Subjective Refraction (UTSR; example in panels B and D). Subjects wore their usual correction while performing the experiments (spectacles or contact lenses) and therefore the result of each repetition (red or blue dots), or the average (solid black line in the center of the gray band indicating the standard deviation), represent the residual refraction over their usual correction.

**Fig. 3.**
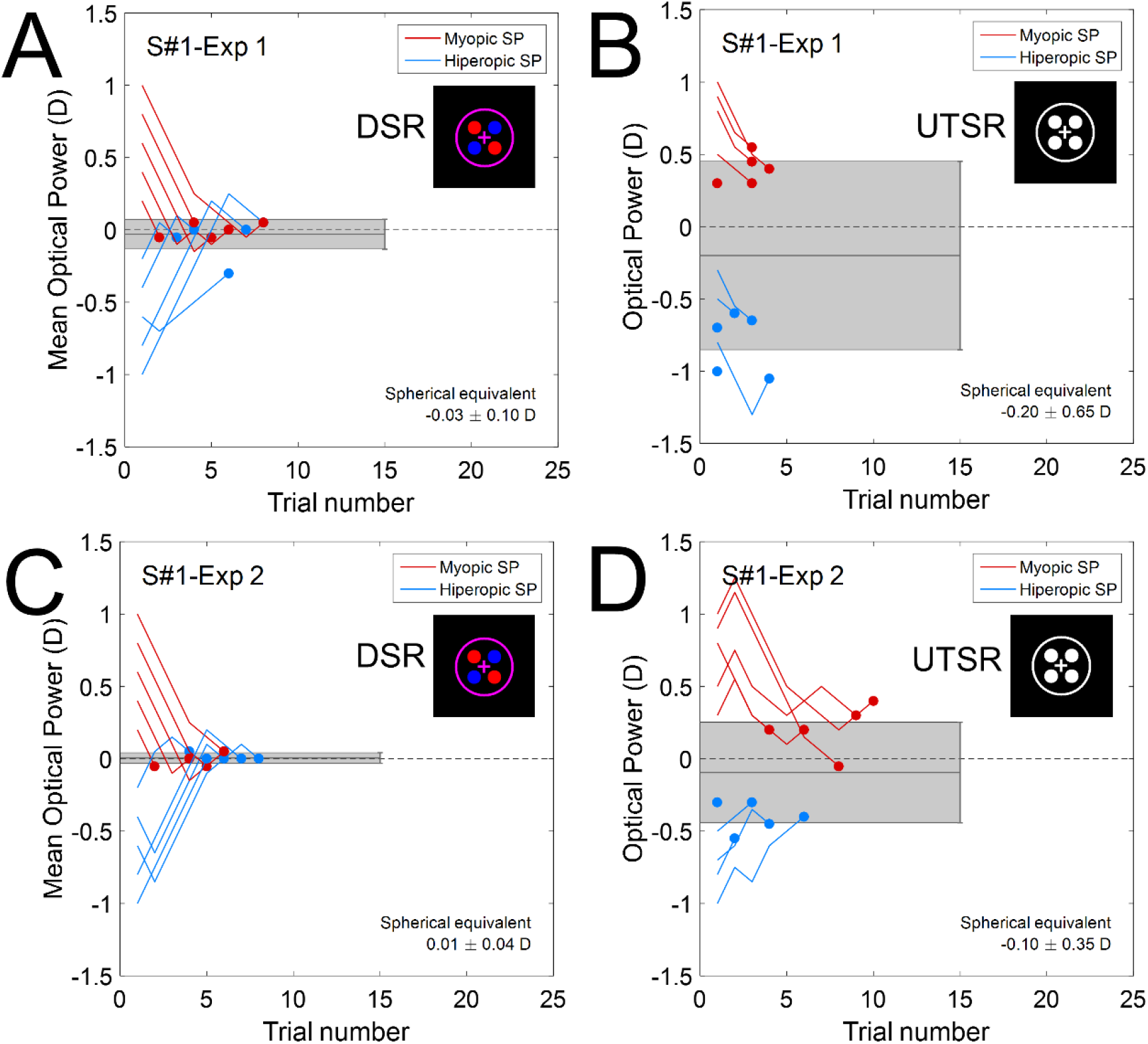
Progress of DSR and UTSR tasks for subject #1 and Experiment 1 and 2. Each panel shows the progress of a subject while performing a visual task (DSR or UTSR) to obtain the subjective refraction. Red lines represent repetitions with myopic starting points and blue lines with hyperopic starting points. The filled dot at the end of each line indicates the residual refraction for that repetition. The gray bar indicates the mean and the standard deviation of the residual refraction across repetitions. These values are indicated in the left-bottom corner of each panel. A miniature of the stimulus used in each example is shown in the upper-right corner. **A**. Evolution of the mean optical power of the TDW (in D) versus the trial number for subject #1 performing the DSR task in Experiment 1. **B**. Optical power (in D) versus the trial number for the same subject (S#1) and experiment while performing the UTSR task. **C**. DSR task in Experiment 2 (paralyzed accommodation) for the same subject (S#1). **D**. UTSR task for the same subject (S#1) also for Experiment 2.

Figure 3A illustrates the DSR task for subject S#1 performed in Experiment 1. All repetitions converge to the best spherical equivalent with a standard deviation of ±0.10D. This value is lower than the finest optical power step available in eyecare clinics (±0.25D). Figure 3B shows the corresponding UTSR task for the same subject and experiment. In this case, there is no convergence, and the standard deviation is much higher ±0.65D, indicating that the unsupervised blur-detection task is dramatically affected by accommodation: hyperopic defocus (blue lines) can be compensated with accommodation and the subject (25 years old) perceives the stimulus instantly sharp. Due to the depth of focus of the eye, myopic defocus is also very soon perceived as sharp. Figure 3C shows the results of the DSR task for the same subject (S#1) for Experiment 2 (cycloplegic drugs). Paralyzing the accommodation results in an even lower standard deviation ±0.04D) with the same spherical equivalent (−0.03D in Experiment 2 vs 0.01D in Experiment 1). This suggests that, at least in this subject, accommodation was barely influencing the outcome of the DSR task in Experiment 1 (where the accommodation was free). Figure 3D shows the results for UTSR task for S#1 and accommodation paralyzed. Now, hyperopic defocus, which was accommodated when accommodation was free (Figure 3B), cannot be compensated. Therefore, the standard deviation of UTSR with cycloplegic drugs is much lower (±0.35D). Further analysis will show the result for all the subjects measured with and without paralyzed accommodation.

The convergence of the subject to a unique result in the DSR task, regardless of the starting point for each repetition (myopic and hyperopic), is consistent across subjects and experiments and proves that the accommodative response, although functional, is not elicited during the DSR task. The quick and abrupt changes in optical power produced by the TDW seem to unfasten the accommodation mechanism from the stimulus. The DSR visual task does not require paying attention to blur and concentrates the attention of the patient in luminance flicker and chromatic distortions. Besides, the task takes place in presence of a dynamic baseline blur that cannot be eliminated, and that seems to make accommodation unproductive. On the contrary, as already mentioned and shown in the examples of Figures 3B and 3D, which are representative of the responses of all subjects, the UTSR task is severely affected by accommodation.

Figure 4 shows the spherical equivalent obtained from the DSR (red) and the UTSR (blue) tasks, for all subjects in Experiment 1 (free accommodation) and Experiment 2 (cyclopegia). The position of the bar indicates the mean across repetitions and the length, one standard deviation at each side of the mean. The average spherical equivalent obtained with the DSR task, across subjects shows a myopic shift of -0.33D. The DSR method detects significant residual refractions in 80% of the subjects (measurements significantly different from zero, the usual correction of the subjects, using a 0.9 significance level, i.e. red bars not touching the zero). UTSR average spherical equivalent has a lower myopic shift (−0.15D) and captures significant residual refractions in only 12% of the subjects.

**Figure 4.**
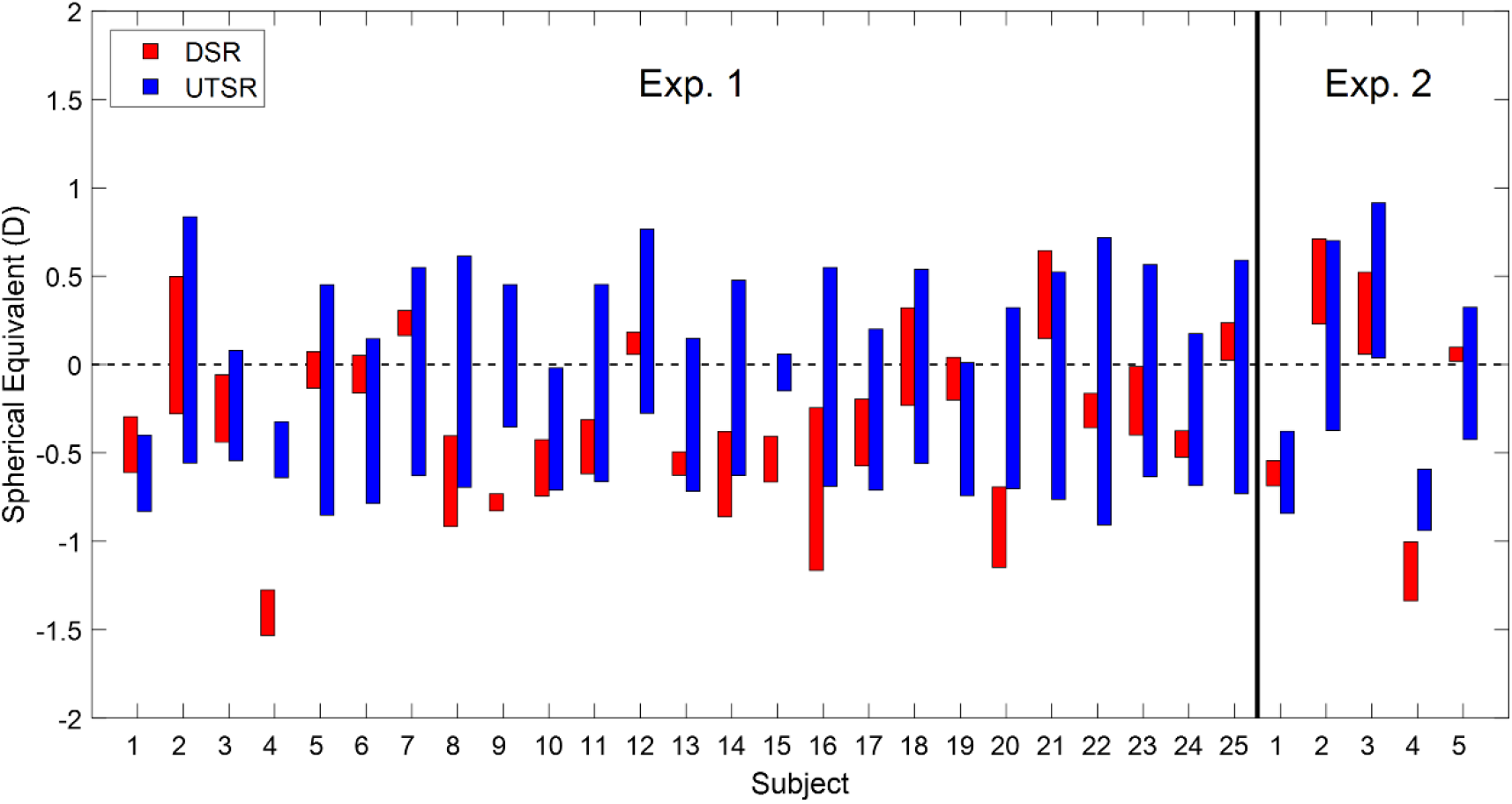
Spherical equivalent and standard deviation for all subjects and experiments. Each bar is centered on the spherical equivalent and its length represents twice standard deviation. Red bars correspond to the DSR task and blue bars to the UTSR task.

Analyzing separately the results of DSR from the two experiments, the average spherical and average standard deviation equivalent across subjects are: - 0.33±0.17D for Experiment 1 and -0.19D±0.15D for Experiment 2. Interestingly, when accommodation is paralyzed (Experiment 2), the average absolute residual error for UTSR (0.41D) is comparable to that found in DSR (0.36D on average across experiments, 0.52D in Experiment 2). For DSR measurements, a Wilcoxon signed-rank test for the same subjects in Experiment 1 vs 2 reports that the differences in mean spherical equivalent were not statistically significant (p=0.19), indicating that, in our sample, paralyzing the accommodation does not induce significant differences.

Figure 5 directly plots the standard deviation across repetitions for each subject and experiment, providing a closer look at the repeatability of the DSR and UTSR tasks. The horizontal dashed lines indicate the average standard deviation across subjects. To compare with the literature, we also plotted the intraoptometrist variability (averaged from [6–8]) with a dark green line and the interoptometrist variability (averaged from [7,9–12,18]) with a light green line. On average, the standard deviation for the DSR task (±0.17D) is 64% lower than that of the UTSR task (±0.47D), 57% lower than the traditional interoptometrist variability (±0.39D), and 39% lower than the intraoptometrist variability (±0.28D). Across the four experiments, the standard deviation for the DSR task is lower than that for the UTSR task in 96.7% of the subjects, lower than the interoptometrist variability in 93.3% of the subjects, and lower than the intraoptometrist variability in 90% of the subjects. These results evidence the higher repeatability (i.e. precision) of the DSR task with respect to the corresponding UTSR task in the same conditions and to the traditional subjective refraction methods. Minimizing flicker and chromatic distortions happens to be more precise than judging blur.

**Figure 5.**
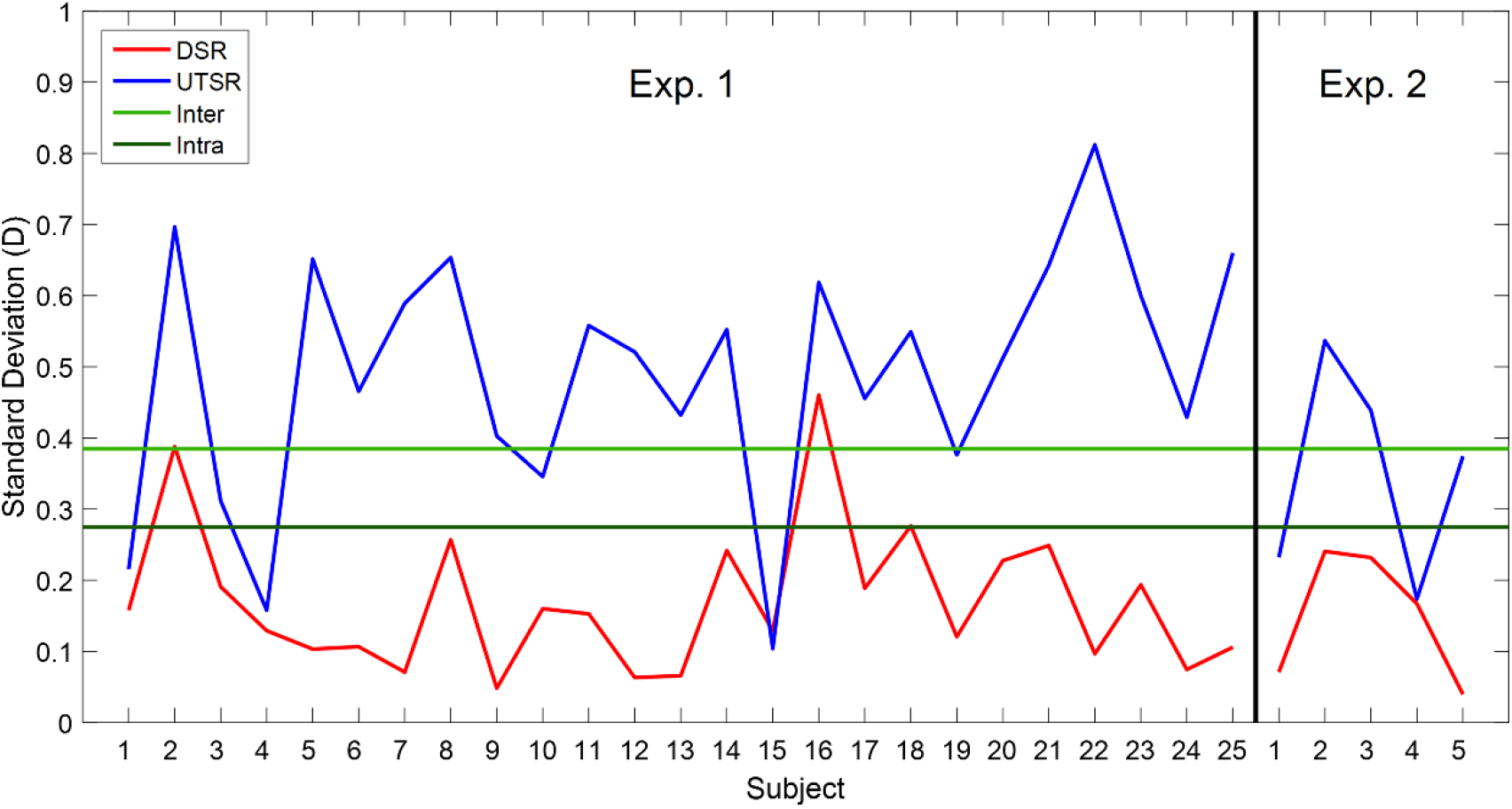
Standard deviation for all subjects and experiments. Standard deviation for the DSR (red curve) and UTSR (blue curve). The horizontal dashed red and blue lines indicate the average standard deviations across subjects for the DSR task and the UTSR task, respectively. The horizontal light green line indicates the interoptometrist variability (traditional subjective refraction) and the dark green light the intraoptometrist variability, both extracted from the literature.

To compare the results obtained with the DSR task and the UTSR task with the Traditional Subjective Refraction (TSR), we performed different Bland-Altman analyses. Figure 6A shows the Bland-Altman plot for the DSR versus the TSR, pooling the data of all repetitions for Experiment 1 (non-cyclopleged subjects). We found a strong correlation (*r*=0.99; *p*<0.001) and an offset of -0.33D. Figure 6B shows the corresponding plot for UTSR (*r*=0.98; *p*<0.001). For UTSR we found similar correlations, and a lower offset (−0.15D), but these results could be artifactual. As we already know from Figures 3B and 3D, the result of each UTSR repetition is strongly affected by the starting point. Since the set of starting points is uniformly distributed within the ±1D range at both sides of the TSR of the subject, the average UTSR outcome could be artificially determined by the average of all starting points and therefore matching the TSR. To explore this potential effect, we performed additional Bland-Altman analyses with subsets of samples in the myopic and hyperopic sides. Figures 6C and 6D correspond to starting points within the +0.8 to +1.0D range (maximum myopia; DSR and UTSR), and Figures 6E and 6F to starting points within the -1.0 to -0.8D range (maximum hyperopia; DSR and UTSR). The correlation coefficients are high in all cases (r=0.99, p<0.001). However, while the DSR offset only changes 0.14D from myopia to hyperopia, the UTSR offset changes as much as 1.23D. The superimposed Limits of Agreement (LoAs, 95% Confidence Interval) along the interval of starting points for DSR are -1.19 to +0.59D (difference 1.78D) and for UTSR -1.53 to +1.20D (difference 2.73D).

**Figure 6.**
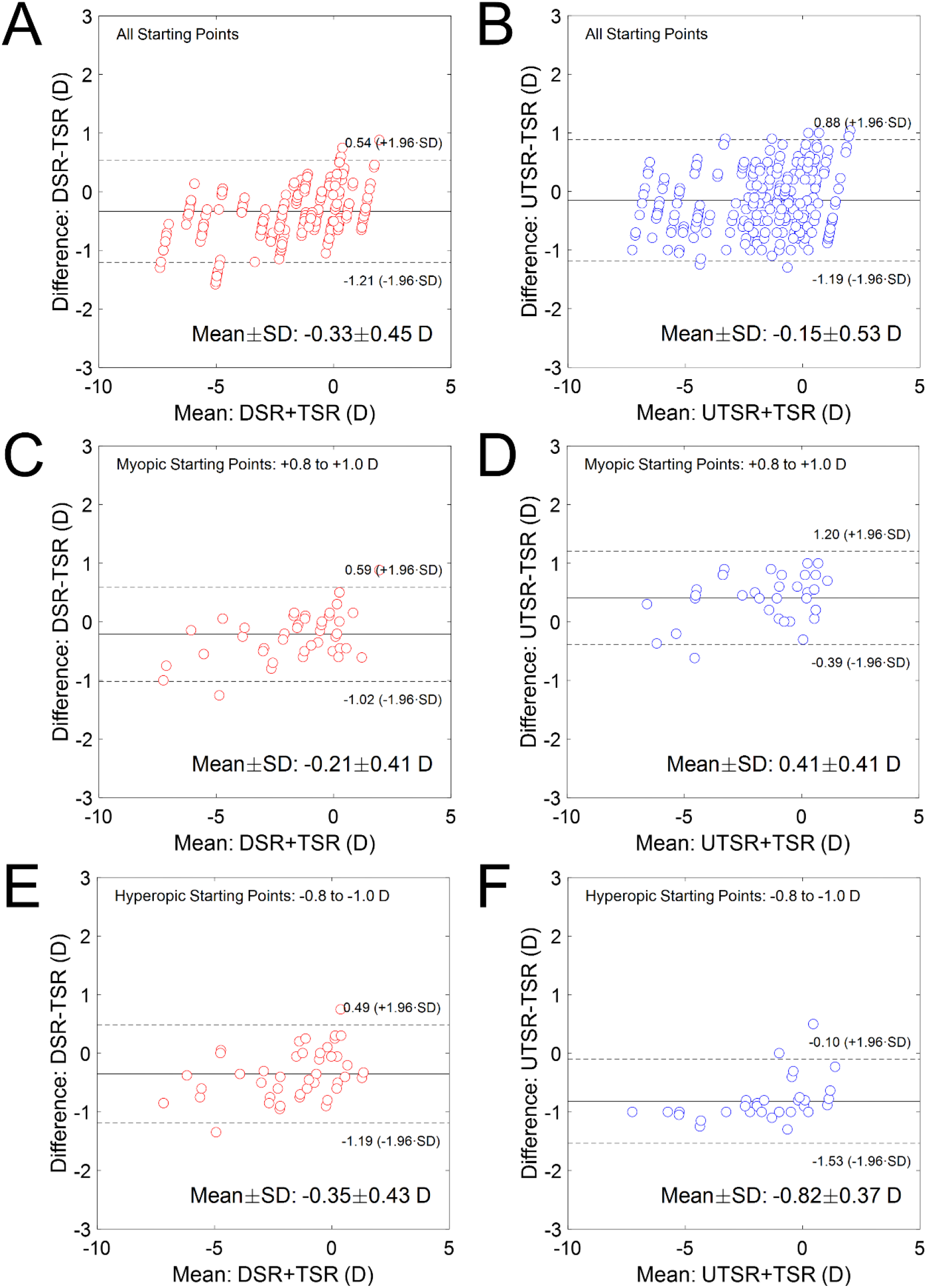
Bland-Altman analysis for Experiment 1. Each panel shows a Bland-Altman plot, indicating the Limits of Agreement (LoA, 95% Confidence Interval), mean, and standard deviation of the sample. **A**. Plot comparing DSR and TSR for all starting points. **B**. Plot comparing UTSR and TSR for all starting points. **C**. Plot comparing DSR and TSR only for highly myopic starting points (+0.8 to +1.0D). **D**. Plot comparing UTSR and TSR only for highly myopic starting points (+0.8 to +1.0D). **E**. Plot comparing DSR and TSR only for highly hyperopic starting points (−1.0 to -0.8D). **F**. Plot comparing UTSR and TSR only for highly hyperopic starting points (−1.0 to -0.8D).

Removing the offset (−0.33D) between DSR and TSR provides a comparison closer to the final implementation of the technique: a similar analysis to Figure 6A, averaging across repetitions and compensating the offset, provides a standard deviation in the residual refraction of ±0.45D and limits of agreement ±0.87D. But the variabilities across subjects and between DSR and TSR reported in this study (standard deviations and LoA) are not only attributable to variability and disparities in the DSR but also include all sources of residual refractive error for TSR (used as baseline): interoptometrist variability, tolerances of glasses and contact lenses, under or overcorrections in the subjects, refraction changes with time, etc. However, the comparison between DSR and UTSR confirms the enormous dependence of the UTSR results on the starting point, due to the well-known detrimental effect of accommodation (in the absence of fogging strategies), that was free and fully functional during Experiment 1 (except for two subjects, S#1 and S#19, that were partially functional due to presbyopia).

The DSR refraction provided higher or similar values of visual acuity, in all the subjects, than the usual refraction of the patients (TSR refraction), but it was not significantly different (paired t-test *p*=0.47).

The spherical equivalent obtained from the TSR and the objective refraction are strongly correlated, and not statistically different (paired t-test *p*=0.32). A Bland-Altman analysis of the objective refraction with respect to the TSR reports a slightly positive deviation of +0.12D and with a mean absolute difference of ±0.54D. The LoAs are -0.93 to +1.17D (difference 2.1D).

Figure 7A explores the effect of accommodation further. The results of the DSR task for all hyperopic starting points (average of all hyperopic repetitions) are directly plotted against the results for all myopic starting points (average of all myopic repetitions), both for DSR (each red symbol indicates a subject) and UTSR (blue symbols). For DSR, the correlation between hyperopic and myopic starting points is very high (*r*=0.94, *p*<0.001) and there is a small but significant statistical difference between them (0.08D on average; paired t-test *p*=0.02). On average, the residual refraction and the standard deviation obtained with myopic starting points is -0.27±0.15D and with hyperopic starting points is -0.35±0.14D. The similarity of the results obtained with hyperopic and myopic starting points with the DSR task demonstrates that this task is barely affected by accommodation and that the method is very precise both for small amounts of myopia and hyperopia. In contrast, for the UTSR method the correlation is low (*r*=0.03, *p*=0.86) between myopic and hyperopic starting points, which provide radically different results (0.17±0.23D and -0.47D±0.30D, average difference 0.47D, paired t-test *p*<0.001). As expected, the unsupervised UTSR method, consisting in a blur-detection task without fogging, with a comparable stimulus and in the same set-up, is severely affected by accommodation, that not only affects repeatability, but also the final spherical equivalent measured, especially in hyperopic patients.

**Figure 7.**
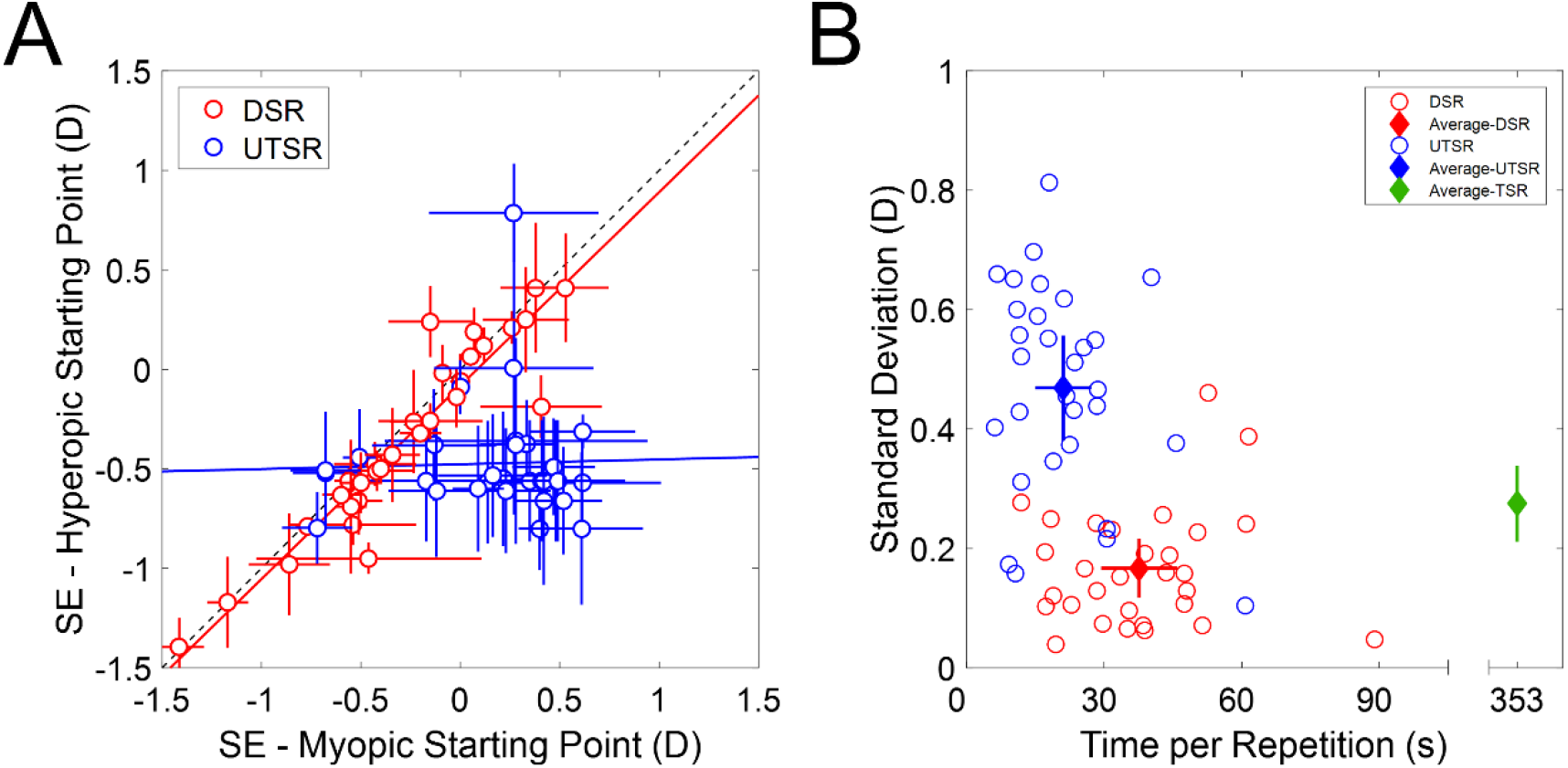
Comparison between DSR and UTSR tasks. **A. Analysis of myopic and hyperopic starting points for DSR and UTSR.** Spherical equivalent obtained from the average of all hyperopic starting points versus the spherical equivalent obtained from the average of all myopic starting points for DSR (red) and UTSR (blue) tasks and all experiments. The error bars indicate the standard deviation across repetitions. **B. Precision and time to perform the tasks**. Standard deviation across repetitions vs the time per repetition for DSR (red), UTSR (blue), and TSR. For TSR, the standard deviation is the intraoptometrist error and the average time is extracted from the literature. The filled diamonds indicate the average across subjects.

Experiment 2 provides additional information about the effect of accommodation. Five subjects carried out the experimental session of Experiment 2 (including the DSR and the UTSR tasks), but under the effect of cycloplegics drugs, paralyzing their accommodation (see *Methods*). Considering the low number of subjects, the correlation between Experiments 1 and 2 (in spherical equivalent) was very high for DSR (*r*=0.92, *p*=0.03), but low and not significant for UTSR (*r*=0.74, *p*=0.15). The slope (ideally 1) was 1.05 for DSR and 1.33 for UTSR). Interestingly, cyclopegia produced a constant shift (y-intercept) of 0.23D in the DSR results, similar and opposite to the average spherical equivalent reported in Experiment 1. The standard deviation was lower for Experiment 2 than for Experiment 1, both for UTSR (on average ±0.35 vs ±0.41D), and DSR (±0.19 vs ±0.15D), but not significant (paired t-tests *p*=0.48 and *p*=0.32).

The ideal method to perform a subjective refraction would provide a good balance between measurement time and variability. Figure 7B shows a scatter plot of the standard deviation in the measurement versus the average time per repetition for all subjects and experiments. We observe two clear clusters: the results for UTSR-blue open circles-have extremely low measurement times but high standard deviations, while the results for DSR -red open circles-have intermediate measurement times and low standard deviations. On average, the UTSR task takes 21±12 seconds per repetition with an average standard deviation of ±0.47±0.18D -solid blue diamond- and the DSR task takes 38±16 seconds per repetition with a standard deviation of ±0.17±0.10D -solid red diamond-. The DSR (also the UTSR) is much faster than the TSR (green diamond), which takes around 350 seconds (almost 6 minutes) according to previous studies [3,5]. DSR is also more precise than TSR (intra-optometrist standard deviation ±0.28D±0.06 [6–8]) and, of course, than UTSR.

Finally, we investigated if the refractive error of the subject influences the DSR method. We divided the subjects into three groups, based on their current correction: hyperopes (>+0.50D; 3 subjects), myopes (<-0.50D; 14 subjects), and emmetropes (±0.50 D; 8 subjects). The results of a Wilcoxon rank-sum test do not show statistically significant differences between the three groups (p>0.05 in all comparisons). Therefore, in our groups of subjects, the refractive error is a statistical variable that does not influence the results.

## Discussion

Subjective refraction is a ubiquitous procedure in eye care clinics. Despite being cumbersome and time-consuming, it has not advanced much in decades. Some technologies as automated phoropters have made the procedure easier but have not improved it. Objective refractors, as wavefront autorefractors, now provide good approximations to subjective refraction but have not been able to replace it [30,31].

Subjective visual tasks are inherently slow, they entail large series of trials, each one requiring a perceptual judgment from the observer (blur detection, blur preference, or letter identification in the case of subjective refraction) and a decision by the practitioner. But the situation is worse in subjective refraction than in other visual tasks. Traditional Subjective Refraction (TSR) begins by displacing the starting point far away from the best estimation available, usually provided by objective refraction (or sometimes by a lensometer), to the myopic side. This long detour in the through-focus trajectory, called fogging, is inefficient in terms of trials, but allows to deal with accommodation, and also provides the direction of focus. An ideal method to measure the subjective refraction would provide a shortcut toward the final spherical equivalent of the patient, using the lowest number of perceptual judgments: only the few steps separating the patient’s subjective focus from the one measured objectively.

In this study we have presented the Direct Subjective Refraction (DSR), a novel technology and methodology to measure the spherical equivalent of the refractive error of an eye, that disentangles, to a large extent, the accommodation mechanism, and that provides the patient with a visual hint of the direction of focus. The starting point can be the best estimation provided by objective refraction, and the spherical equivalent can be found directly. The number of trials (perceptual judgments) is reduced, and each one is straightforward and not supervised, producing faster measurement times than the TSR. The DSR task is direct in the sense that color provides an unambiguous cue for the direction of the next step in the staircase (color distortions are different at both sides of the retina: blueish in the myopic side, and reddish in the hyperopic side).

In the last steps of the TSR, the practitioner checks if power changes of ±0.25D improve the visual acuity or visual comfort, sometimes using colored backgrounds (such as the duochrome test). These final checks inspired the development of the DSR method. Actually, the DSR performs 15 of those optical power changes per second, during all the duration of the measurement and in every step of the subjective staircase leading to the spherical equivalent. Besides, 15 Hz is a frequency providing maximum temporal sensitivity to flicker [42], and therefore optimal for the task, although out of reach for the accommodative system. Hence, DSR provides a much stronger perceptual cue (in fact two concurrent and reinforcing signals: flicker and color) than TSR and is isolated from accommodation (because blur, the main clue for the accommodative system is no longer involved in the task), allowing straightforward measurements for the patients without requiring the guidance of the clinician. Our results show that DSR provides a precise, accurate, and fast estimation of the spherical equivalent of the subjective refraction.

To put the results of the DSR task in the appropriate context, we have confronted them with comparable data of TSR obtained from the literature, and with the UTSR (Unsupervised Traditional Subjective Refraction) task. UTSR is a quick version of TSR, also depending on blur judgments but unsupervised and without fogging techniques. UTSR can also be considered a black-and-white and static version of the DSR.

As seen in Figures 3-5, UTSR has more variability in the spherical equivalent (standard deviation ±0.47D) than TSR (±0.28D). At the same time, with DSR we found (Figure 5A) subclinical variability across repetitions in the spherical equivalent (±0.17D) and an average offset -0.33D with respect to the correction worn by the subjects. This offset, although small, is clinically relevant. However, it has minor importance because it represents a systematic shift in the measurements (as discussed below) that could ultimately be compensated with a correction factor.

Accommodation is a potential cause of variability and systematic shifts during any subjective refraction technique (also during objective refraction). To study the effect of accommodation in the UTSR and DSR tasks, we included different starting points in the different repetitions, simulating different amounts of myopia or hyperopia in the same subject. We found that UTSR is undoubtedly affected by dynamic accommodation. In contrast to the TSR procedures used in clinics, that implement different strategies to reduce the impact of accommodation and to reach the center of the depth of focus interval, UTSR is not protected against accommodation and results in important variabilities due to offsets that depend on the starting point: between +0.41D if the starting point is myopic, and -0.82D if hyperopic (Figure 6). Hyperopic starting points can be accommodated (red points in Figure 3A and Figure 3C), bringing the image into focus and finishing the staircase prematurely, leaving a negative offset (underestimating the correction). Similarly, depth of focus produces positive shifts in myopic defocus (overestimating the correction) because subjects judge the image sharp before reaching the maximum optical quality, and stop the staircase fractions of diopter in front of the best focus (blue points in Figure 3B and Figure 3D). A direct comparison of the different starting points simulating myopia or hyperopia (Figure 6, Figure 7A) reinforces the finding that DSR is unaffected by the accommodative response, which spoils the UTSR measurements.

As seen in the examples of Figure 3, the DSR task forces the subject to reach the best focus and the staircases oscillate at both sides of it, removing the positive offset associated with myopic starting points. Figure 7A suggests that, at the same time, the accommodation mechanism is to a large extent deactivated during the DSR measurements. Defocus is present in the stimulus during DSR, as in UTSR, and is certainly perceived by the subject, but it is not part of the DSR task. The accommodation of the eye remains in a fixed state because the fast change induced by the TDW prevents its activation. The fast defocus alternation in the image does not provide a fixed reference to focus and does not elicit accommodation. However, the small offset found could still be attributed to a small remaining residual accommodation (tonic accommodation). Being stable and largely unaffected by the stimulus, we could refer to this accommodative state as the ‘resting position’ of the eye, in a ‘dark focus’ of ‘tonic accommodation’ closer than infinity, previously reported in conditions where the accommodation is lost, for example in night myopia [43–45].

Experiment 2 corroborated these findings. Results before and after cyclopegia provided insignificant correlations with UTSR (and slope 1.33) but were highly correlated for DSR (and with slope 1.05). Paralyzing accommodation also had the effect of reducing the measurement variability to its lowest value (±0.15D on average; in DSR). Although promising, these findings should be confirmed in a higher number of patients.

Accommodation to the stimulus can be discarded as an explanation to the offset found in DSR, but several other reasons could provide a plausible explanation. For example, DSR contains the implicit assumption that the spherical equivalent lays in the intermediate position (in diopters) between the red focus and the blue focus (Figure 1). Therefore, changes in the spectral composition of the stimulus (spectral width and position of the red and blue peaks) could shift the spherical equivalent measured with DSR. Besides, the relationship between wavelength and focus position in diopters (the LCA curve) is not lineal [46,47], and the monochromatic wavelength-in-focus for subjective refraction changes with the subject [48], making the prediction on the polychromatic spherical equivalent from monochromatic measurements difficult to model [49]. Furthermore, even the gold standard, the polychromatic spherical equivalent measured with TSR, can change with the color temperature of the white light used. Moreover, the variability of any subjective measurement (TSR, DSR, UTSR) is extremely related to the subjective depth of focus, not only for optical reasons (aberrations) but for neuronal reasons [50].

Instrument myopia (an effect also related to accommodation) or pupil size effects due to relatively low ambient light levels during the measurements, such as potential focus shifts due to spherical aberration or to depth of focus increments, could also be blamed for this small but significant offset between DSR and TSR. In fact, the magnitude of most of the mentioned effects, separately, could be higher than the offset found in our measurements [48–50]. The calibration of the instrument and the fidelity of the TDW to the nominal power, as well as the distances involved in the optical set-up, were checked before the measurements, and the potential deviations in optical power could be considered negligible [51]. Still, further research of all these hypotheses under clinical conditions in a large number of patients will allow the development of strategies to null or compensate the offset.

The fogging techniques required to reduce the influence of accommodation make TSR tiresome for the subject and the practitioner and increase the measurement time. TSR is indeed a time-consuming procedure, reported taking almost 6 minutes per subject, on average [3–5]. An unsupervised version of the traditional procedure, the UTSR, is in fact very quick, taking about 21 seconds per repetition. However, it has low precision, with repeatability across subjects of ±0.47D, and systematic deviations that depend on the starting point. The DSR method, insensitive to accommodation by design, does not require fogging strategies. It is not only a very precise (±0.17D) and accurate procedure (−0.33D without offset compensation), but also fast: it takes less than 40 seconds to be performed, on average. The fastest subject took only 12 seconds per repetition, and the slowest, 90 seconds. The short measurement time (plus only 1.5 minutes of explanation) allows thorough training and would allow several repetitions, although only a few are really needed (probably two, one for training and approximation and another one for refinement), given the high repeatability of the method. An assessment of reliability of Experiment 1 supports these conclusions, as it reports a Cronbach’s Alpha value of 0.979 -with reliability factor 0.95-estimates that only the first 4 repetitions are not redundant.

In this study, we measured the spherical equivalent, but not the amount of astigmatism in the subjective correction. Moreover, we measured the subjects with their usual corrections on, therefore reducing the amount of astigmatism, if present, to a residual value. Pilot experiments have showed that the DSR task can be performed in presence of astigmatism [52], at least up to one diopter bur presumably more. This promising result suggests that the DSR approach described could also have potential for fast and unsupervised measurement of the subjective refraction including astigmatism. For that, the current method should be refined with new stimuli, oriented features, and new measurement protocols considering flicker and chromatic distortions in different orientations.

Some autorefractors have achieved comparable repeatability in non-cyclopeged eyes as the one reported here with DSR, and are evolving into hand-held and binocular instruments [53]. But DSR is a subjective method to measure the spherical equivalent, and as such with true potential to replace TSR with a faster and direct approach. The DSR technology is as simple as projecting an optotunable lens onto the eye and can be easily implemented in portable binocular devices, hand-held or even wearable [40,54,55], potentially substituting phoropters or trial frames in clinical centers or in screening campaigns outside the clinic.

## CONCLUSIONS

The Direct Subjective Refraction presented here is a straightforward and unsupervised new subjective method to obtain the spherical equivalent of an eye. Direct Subjective Refraction overpasses existing subjective methods in terms of accuracy, precision, and measurement time, with the potential to become a new paradigm in the measurement of subjective refraction.

## Acknowdlegments

The project that gave rise to these results received the support of a fellowship from “la Caixa” Foundation (ID 100010434; LCF/BQ/DR19/11740032) and Spanish Ministry of Education Grant FPU17/02760 to V.R.L., and Spanish Ministry of Science, Innovation, and Universities Grants ISCIII DTS2016/00127, FIS2017-84753-R and CSIC LINKA20122 to C.D.

## Author contributions

V.R.L. collected and analyzed data, A.H.P. collected data; V.R.L., and C.D. conceived the project and wrote and edited the paper.

